# AutoVEM2: a flexible automated tool to analyze candidate key mutations and epidemic trends for virus

**DOI:** 10.1101/2021.05.08.443047

**Authors:** Binbin Xi, Shuhua Li, Wei Liu, Dawei Jiang, Yunmeng Bai, Yimo Qu, Jerome Rumdon Lon, Lizhen Huang, Hongli Du

## Abstract

In our previous work, we developed an automated tool, AutoVEM, for real-time monitoring the candidate key mutations and epidemic trends of SARS-CoV-2. In this research, we further developed AutoVEM into AutoVEM2. AutoVEM2 is composed of three modules, including call module, analysis module, and plot module, which can be used modularly or as a whole for any virus, as long as the corresponding reference genome is provided. Therefore, it’s much more flexible than AutoVEM. Here, we analyzed three existing viruses by AutoVEM2, including SARS-CoV-2, HBV and HPV-16, to show the functions, effectiveness and flexibility of AutoVEM2. We found that the N501Y locus was almost completely linked to the other 16 loci in SARS-CoV-2 genomes from the UK and Europe. Among the 17 loci, 5 loci were on the S protein and all of the five mutations cause amino acid changes, which may influence the epidemic traits of SARS-CoV-2. And some candidate key mutations of HBV and HPV-16, including T350G of HPV-16 and C659T of HBV, were detected. In brief, we developed a flexible automated tool to analyze candidate key mutations and epidemic trends for any virus, which would become a standard process for virus analysis based on genome sequences in the future.

**Highlights:**

1. An automatic tool to quickly analyze candidate key mutations and epidemic trends for any virus was developed.
2. Our integrated analysis method and tool could become a standard process for virus mutation and epidemic trend analysis based on genome sequences in the future.
3. N501Y with the other 16 highly linked mutation sites of SARS-CoV-2 in the UK and Europe were further confirmed, and some valuable mutation sites of HBV and HPV-16 were detected.

## 1. Introduction

SARS-CoV-2 has infected over 151,812,556 people and caused 3,186,817 deaths by 2 May 2021 [1]. At present, a variety of vaccines against SARS-CoV-2 are being used over the world, including mRNA-1273 [2], BNT162b2 [3], CoronaVac [4] and so on, hoping to form the effect of herd immunity. However, it is reported that N501Y mutation in the spike protein may reduce the neutralization sensitivity of antibodies, and finally influence the effectiveness of some vaccines [5]. Therefore, real-time monitoring the epidemic trend of SARS-CoV-2 mutations is of great significance to the update of detection reagents and vaccines. In our previous work, we found 9 candidate key mutations [6], including A23403G causing D614G amino acid change on the S protein, which has been proved to increase the infectivity of SARS-CoV-2 by several in vitro experiences [7-11]. With the further global spread of SARS-CoV-2, it is difficult to prevent its mutation. Therefore, we proposed an innovative and integrative method that combines high-frequency mutation site screening, linkage analysis, haplotype typing and haplotype epidemic trend analysis to monitor the evolution of SARS-CoV-2 in real time. And we developed the whole process into an automated tool: AutoVEM [12]. We further found that the 4 highly linked sites (C241T, C3037T, C14408T and A23403G) of the previous 9 candidate key mutations have been almost fixed in the virus population, and the other 5 mutations disappeared gradually [12]. In addition, we found another 6 candidate key mutations with increased frequencies over time [12].

Our research on the trend of haplotype prevalence and other studies on the trend of single site prevalence both show that SARS-CoV-2 is constantly emerging new mutations, and the frequency of some mutations is increasing over time, while the frequency of some mutations is decreasing or even completely disappearing over time [6, 12, 13]. The consistent findings indicated that the integrative method we proposed is reliable. Moreover, the haplotype prevalence trend we used makes the new epidemic mutants less complicated. However, AutoVEM we developed is only for SARS-CoV-2 analysis. With the changes in the global natural environment, new and sudden infectious diseases are continuously emerging, such as the outbreak of SARS in Feb 2003 [14], MERS in 2012 [15], Ebola in 2014 [16], and the ZIKV in 2015 [17]. Therefore, we need a more flexible automated tool to identify and monitor the key mutation sites and evolution of various viruses.

In this research, we further developed AutoVEM into AutoVEM2. AutoVEM2 is composed of three different modules, including call module, analysis module and plot module. The call module can carry out quality control of genomes and find all SNVs for any virus genome sequences with various optional parameters. The analysis module can carry out candidate key mutations screening, linkage analysis, haplotype typing with optional parameters of mutation frequency and mutation sites. And the plot module can visualize the epidemic trends of haplotypes. The three modules can be used modularly or as a whole for any virus, as long as the corresponding reference genome is provided. Therefore, AutoVEM2 is much more flexible than AutoVEM. Here, we analyzed 3 existing viruses by AutoVEM2, including SARS-CoV-2, HBV and HPV-16, to show its functions, effectiveness and flexibility. The SARS-CoV-2 genomes from the UK, Europe, and the USA were analyzed separately due to their large number of SARS-CoV-2 genomes in the GISAID. In addition to existing viruses, AutoVEM2 can also be used to analyze any virus that may appear in the future. We think our integrated analysis method and tool could become a standard process for virus mutation and epidemic trend analysis based on genome sequences in the future.

## 2. Materials and Methods

### 2.1. Functions of Three Modules of AutoVEM2

AutoVEM2 is a highly specialized, flexible, and modular pipeline for quickly monitoring the candidate key mutations, haplotype subgroups, and epidemic trends of different viruses by using virus whole genome sequences. It is written in Python language, in which Bowtie 2 [18], SAMtools [19], BCFtools [20], VCFtools [21] and Haploview [22] are used. AutoVEM2 consists of three modules, including call module, analysis module, and plot module, which can be used modularly or as a whole, and each module performs specific function(s) (Fig 1).

**Figure 1:**
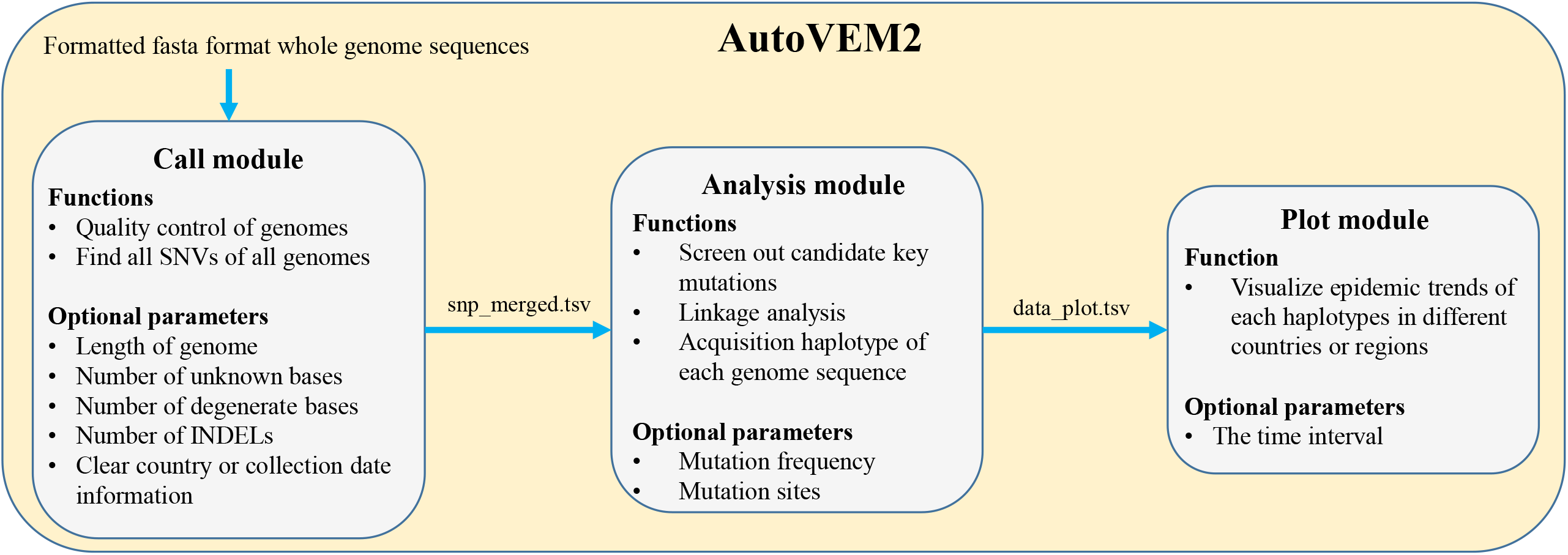
Functions and optional parameters of three modules of AutoVEM2

#### 2.1.1. Call Module

The call module performs the function of finding all SNVs for all genome sequences. The input of the call module is a folder that stores formatted fasta format genome sequences. The call module processes are as follows:

1. Quality control of genome sequence according to four optional parameters: --length, --number_n, --number_db, and --region_date_filter.
2. Align the genome sequence to the corresponding reference sequence by Bowtie 2 v2.4.2 [18].
3. Call SNVs and INDELs by SAMtools v1.10 [19] and BCFtools v1.10.2 [20], resulting in a file named Variant Call Format (VCF) containing both SNVs and INDELs information.
4. Further quality control of genome sequence according to the --number_indels optional parameter. Remove all INDELs from the sequence that has passed further quality control by VCFtools v0.1.16 [21], resulting in a VCF file that only contains SNVs information.
5. Merge SNVs for all genome sequences, resulting in a Tab-Separated Values (tsv) file named snp_merged.tsv.

#### 2.1.2. Analysis Module

The analysis module performs three functions: screening out candidate key mutations, linkage analysis of these candidate key mutations, and acquisition haplotype of each genome sequence according to the result of linkage analysis. The input is the snp_merged.tsv file produced by the call module. The analysis module processes are as follows:

1. Count the mutation frequency of all mutation sites.
2. Screen out candidate key mutation sites according to the --frequency (default 0.05) optional parameter, and candidate key mutation sites can also be specified by the --sites optional parameter.
3. Nucleotides at these specific sites of each genome are extracted and organized according to the order of genome position.
4. Linkage analysis of these specific sites by Haploview v4.2 [22].
5. Acquire haplotypes using Haploview v4.2 [22]. Define the haplotype of each genome sequence according to the haplotype sequence, and if frequency of one haplotype <1%, it will be defined as “other”. This finally results in a tsv file named data_plot.tsv.

#### 2.1.3. Plot Module

The plot module performs the function of visualizing epidemic trends of each haplotype in different countries or regions. The input of the plot module is the data_plot.tsv file produced by the analysis module. The plot module processes are as follows:

1. Divide the whole time into different time periods according to the --days parameter.
2. Count the number of different haplotypes in each time period of different countries or regions.
3. Visualize the statistical results.

### 2.2. Genome Sequences Acquisition, Pre-processing, and Analyzing

SARS-CoV-2 whole genome sequences of the United Kingdom, Europe, and the United States were downloaded from GISAID between 01 Dec 2020 and 28 Feb 2021, resulting in 93,262, 161,703, and 40,405 genome sequences, respectively (Table 1). All HBV and HPV-16 nucleotide sequences, including whole genome sequences and fragments of whole genome, were downloaded from NCBI, resulting in 119,721 and 10,269 sequences, respectively (Table 1). Reference genome sequences of the three viruses were downloaded from NCBI (Table 1). The genome sequences were processed by in-house python script to make them meet the input format of AutoVEM2. Each formatted sequence consisted of two sections, the head section and the body section. The head section started with a greater than sign, followed by the virus name, sequence unique identifier, sequence collection time, and country or region where the sequence was collected, which were separated by vertical lines. And the body section was the nucleotide sequence.

**Table 1.** Information of SARS-CoV-2, HBV, and HPV-16 genomes and the analysis process of the three viruses

| Virus | Sequences<br>Collection<br>Date | Database | Number of<br>Downloaded<br>Genomes | Number of<br>Filtered<br>Genomes<sup>1</sup> | Reference<br>Sequence | Find<br>all<br>SNVs | Screen Out<br>Candidate Key<br>Mutation Sites | Linkage<br>Analysis and<br>Acquire<br>Haplotypes | Epidemic<br>Trends of<br>Haplotypes |
| --- | --- | --- | --- | --- | --- | --- | --- | --- | --- |
| SARS-CoV-2<br>(UK) | 2020.12.01 -<br>2021.02.28 | GISAID | 93,262 | 79,269 | NC_045512.2 | yes | yes | yes | yes |
| SARS-CoV-2<br>(Europe) | 2020.12.01 -<br>2021.02.28 | GISAID | 161,703 | 139,703 | NC_045512.2 | yes | yes | yes | yes |
| SARS-CoV-2<br>(USA) | 2020.12.01 -<br>2021.02.28 | GISAID | 40,405 | 30,142 | NC_045512.2 | yes | yes | yes | yes |
| HBV | - 2021.01.25 | NCBI | 119,721 | 11,088 | NC_003977.2 | yes | yes | yes | no |
| HPV-16 | - 2021.01.25 | NCBI | 10,269 | 1,637 | K02718 | yes | yes | yes | no |

<sup>1</sup> Filtered criteria: the genomic sequences with more than 90% full length and less than 1% N were retained for HBV and HPV-6, while the filtered criteria for SARS-CoV-2 genome was referred to AutoVEM<sup>[12]</sup>

For SARS-CoV-2, sequences with length < 29000, number of unknown bases > 15, number of degenerate bases > 50, number of indels > 2, or unclear collection time information or country information were filtered out [6, 12]. Finally, there were 79,269 sequences of the UK, 139,703 sequences of Europe, and 30,142 sequences of the USA (Table 1). All SNVs of these genomes were found by the call module. Mutation sites with mutation frequency ≥ 0.15 of the UK and Europe (in order to include the five high linkage sites we found before [12]), and 0.25 of the USA would be as their candidate key mutation sites. Linkage analysis of these specific sites was performed and haplotype of each genome sequence was obtained by the analysis module. Epidemic trends of each haplotype were visualized by the plot module. (Table 1)

For HBV and HPV-16, sequences with length <90% and the number of unknown bases >1% the length of reference genomes were filtered out, resulting in 11,088 HBV genome sequences and 1,637 HPV-16 genome sequences. All SNVs of HBV and HPV-16 were found using the call module. Mutation sites with mutation frequency ≥ 0.25 of HBV and HPV-16 would be as the candidate key mutations. Linkage analysis of these specific sites was performed and haplotype of each genome sequence was obtained by the analysis module. (Table 1)

### 2.3. Variation Annotation

The candidate key mutation sites of SARS-CoV-2 in the UK, Europe, and the USA were annotated by an online tool of China National Center for Bioinformation (https://bigd.big.ac.cn/ncov/online/tool/annotation?lang=en), respectively. The candidate key mutation sites of HBV and HPV-16 were annotated manually.

## 3. Results

### 3.1. Overview of the analysis results of SARS-CoV-2, HBV, and HPV-16

The same 27 candidate key mutation sites were screened from the 79,269 SARS-CoV-2 (UK) and 139,703 SARS-CoV-2(Europe) genomes. Through linkage analysis of the 27 sites, it can be divided into 6 and 5 haplotypes with a proportion ≥1% for the UK and Europe, respectively. 13 candidate key mutation sites were screened from the 30,142 SARS-CoV-2(USA) genomes. Through linkage analysis of the 13 sites, the SARS-CoV-2 in the USA can be divided into 21 haplotypes with a proportion ≥1%. (Table 2) 7 of HBV and 12 of HPV-16 candidate key mutation sites were found from the 11,088 HBV genomes and 1,637 HPV-16 genomes, respectively. HBV and HPV-16 can be divided into 24 and 18 haplotypes with a proportion ≥1% by the 7 sites and 12 sites, respectively. (Table 2)

**Table 2.** Candidate key mutation sites and haplotypes results of SARS-CoV-2, HBV, and HPV-16

| Virus | Number of<br>Candidate<br>Key<br>Mutation<br>Sites | Candidate Key Mutation Sites | Number of<br>Haplotypes<sup>1</sup> |
| --- | --- | --- | --- |
| SARS-CoV-2 (UK) | 27 | C241T T445C C913T C3037T C3267T<br>C5388A C5986T C6286T T6954C<br>C14408T C14676T C15279T T16176C<br>A17615G C22227T A23063T A23403G<br>C23604A C23709T T24506G G24914C<br>C26801G C27972T G28048T A28111G<br>C28977T G29645T | 6 |
| SARS-CoV-2<br>(Europe) |  |  | 5 |
| SARS-CoV-2<br>(USA) | 13 | C241T C1059T C3037T C10319T<br>C14408T A18424G C21304T A23403G<br>G25563T G25907T C27964T C28472T<br>C28869T | 21 |
| HBV | 7 | T192G T456C C659T C669T A1546T<br>G2337A G2479A | 24 |
| HPV-16 | 12 | T350G A2925G C3409T A3977C<br>A4040G T4226C A4363T G4936A<br>A5224C C6240G A6432G G7191T | 18 |

<sup>1</sup>Haplotypes with a proportion $\geq 1\%$

### 3.2. Linkage and haplotype analysis of the 27 sites with a frequency ≥0.15 of SARS-CoV-2 in the United Kingdom and Europe

The detailed information for the 27 candidate key mutation sites screened from the UK and Europe was showed in Table 3. According to the linkage analysis, only 6 and 5 haplotypes with a frequency ≥1% were found and accounted for 93.47% and 85.77% of SARS-CoV-2 population in the UK and Europe, respectively (Table 4), which showed highly linked among the 27 candidate key mutation sites (Fig 2A, Fig 2B).

**Table 3.** The annotation of the 27 sites of SARS-CoV-2(UK and Europe) with a mutation frequency ≥15%

| Position | Ref | Alt | Frequency<br>UK<sup>1</sup> | Frequency<br>Europe<sup>2</sup> | Gene Region | Mutation<br>Type | Protein<br>Changed | Codon Changed | Impact |
| --- | --- | --- | --- | --- | --- | --- | --- | --- | --- |
| 241 | C | T | 0.9659 | 0.9552 | 5'UTR | upstream | NA | NA | MODIFIER |
| 445 | T | C | 0.1801 | 0.2394 | gene-orf1ab | synonymous | 60V | 180gtT>gtC | LOW |
| 913 | C | T | 0.7831 | 0.5692 | gene-orf1ab | synonymous | 216S | 648tcC>tcT | LOW |
| 3,037 | C | T | 0.9779 | 0.9737 | gene-orf1ab | synonymous | 924F | 2772ttC>ttT | LOW |
| 3,267 | C | T | 0.7892 | 0.5794 | gene-orf1ab | missense | 1001T>I | 3002aCt>aTt | MODERATE |
| 5,388 | C | A | 0.7881 | 0.5738 | gene-orf1ab | missense | 1708A>D | 5123gCt>gAt | MODERATE |
| 5,986 | C | T | 0.7891 | 0.5827 | gene-orf1ab | synonymous | 1907F | 5721ttC>ttT | LOW |
| 6,286 | C | T | 0.1813 | 0.2421 | gene-orf1ab | synonymous | 2007T | 6021acC>acT | LOW |
| 6,954 | T | C | 0.7896 | 0.5799 | gene-orf1ab | missense | 2230I>T | 6689aTa>aCa | MODERATE |
| 14,408 | C | T | 0.9718 | 0.9679 | gene-orf1ab | missense | 4715P>L | 14144cCt>cTt | MODERATE |
| 14,676 | C | T | 0.7862 | 0.5747 | gene-orf1ab | synonymous | 4804P | 14412ccC>ccT | LOW |
| 15,279 | C | T | 0.7904 | 0.5801 | gene-orf1ab | synonymous | 5005H | 15015caC>caT | LOW |
| 16,176 | T | C | 0.7862 | 0.5745 | gene-orf1ab | synonymous | 5304T | 15912acT>acC | LOW |
| 17,615 | A | G | 0.2579 | 0.1790 | gene-orf1ab | missense | 5784K>R | 17351aAg>aGg | MODERATE |
| 22,227 | C | T | 0.1810 | 0.2439 | gene-S | missense | 222A>V | 665gCt>gTt | MODERATE |
| 23,063 | A | T | 0.7860 | 0.5777 | gene-S | missense | 501N>Y | 1501Aat>Tat | MODERATE |
| 23,403 | A | G | 0.9914 | 0.9770 | gene-S | missense | 614D>G | 1841gAt>gGt | MODERATE |
| 23,604 | C | A | 0.7913 | 0.5829 | gene-S | missense | 681P>H | 2042cCt>cAt | MODERATE |
| 23,709 | C | T | 0.7854 | 0.5748 | gene-S | missense | 716T>I | 2147aCa>aTa | MODERATE |
| 24,506 | T | G | 0.7858 | 0.5740 | gene-S | missense | 982S>A | 2944Tca>Gca | MODERATE |
| 24,914 | G | C | 0.7847 | 0.5739 | gene-S | missense | 1118D>H | 3352Gac>Cac | MODERATE |
| 26,801 | C | G | 0.1739 | 0.2365 | gene-M | synonymous | 93L | 279ctC>ctG | LOW |
| 27,972 | C | T | 0.7725 | 0.5630 | gene-ORF8 | stop | 27Q>* | 79Caa>Taa | HIGH |
| 28,048 | G | T | 0.7862 | 0.5695 | gene-ORF8 | missense | 52R>I | 155aGa>aTa | MODERATE |
| 28,111 | A | G | 0.7834 | 0.5716 | gene-ORF8 | missense | 73Y>C | 218tAc>tGc | MODERATE |
| 28,977 | C | T | 0.7746 | 0.5690 | gene-N | missense | 235S>F | 704tCt>tTt | MODERATE |
| 29,645 | G | T | 0.1807 | 0.2370 | gene-ORF10 | missense | 30V>L | 88Gta>Tta | MODERATE |

<sup>1</sup>Mutation frequency of the 27 sites of 79,269 SARS-CoV-2 genomes from the UK
<sup>2</sup>Mutation frequency of the 27 sites of 139,703 SARS-CoV-2 genomes from Europe

**Table 4.**
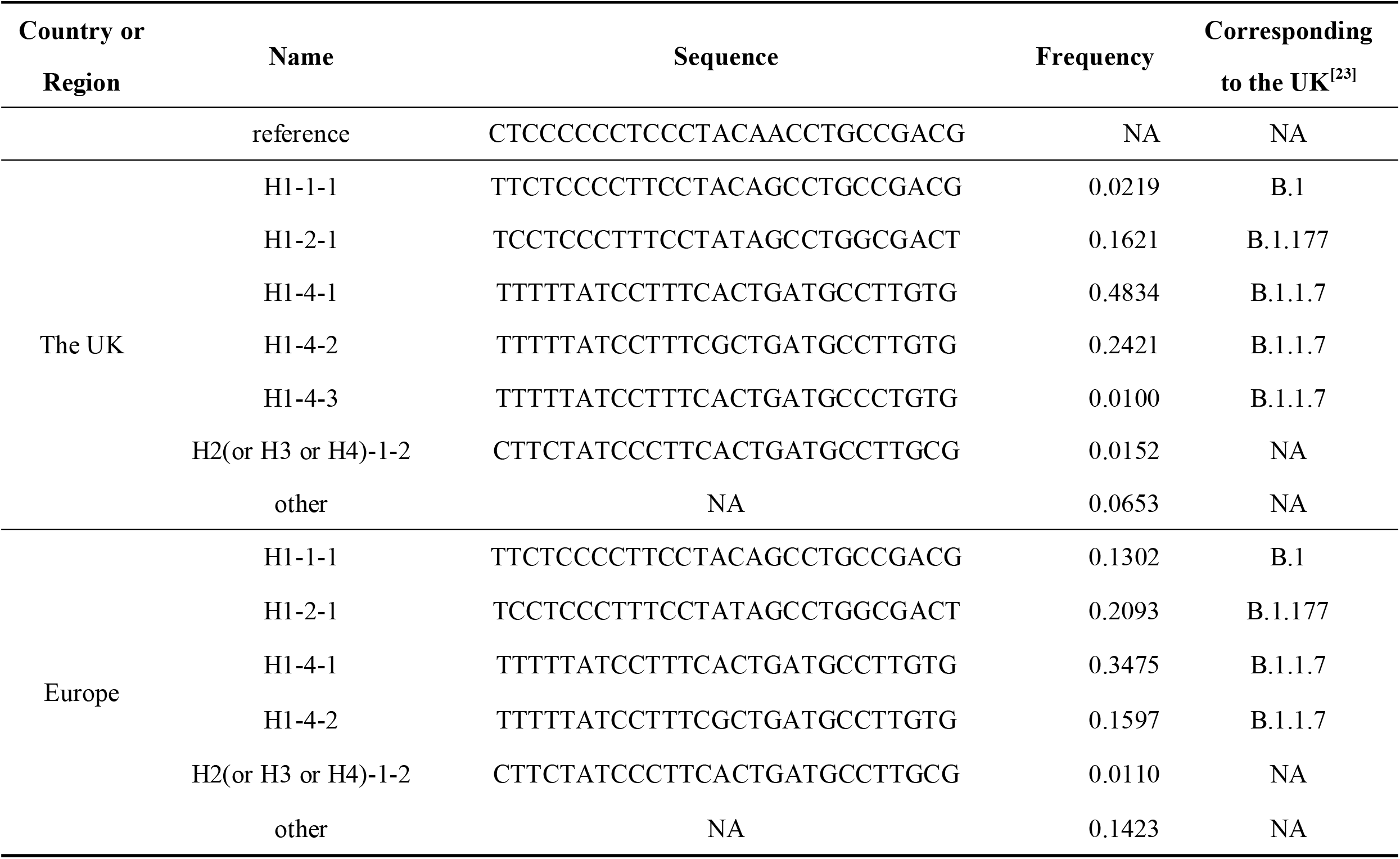
Haplotypes and their frequencies of the 27 sites of SARS-CoV-2(UK and Europe)

**Figure 2:**
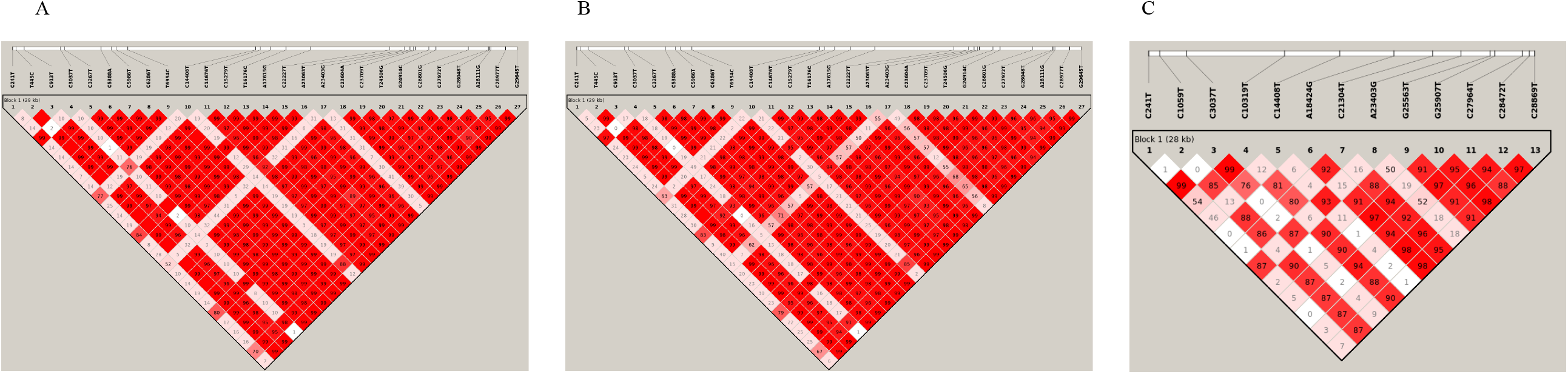
Linkage analysis results of SARS-CoV-2. A. Linkage analysis of 27 candidate key mutation sites of SARS-CoV-2 (UK); B. Linkage analysis of 27 candidate key mutation sites of SARS-CoV-2 (Europe); C. Linkage analysis of 13 candidate key mutation sites of SARS-CoV-2 (USA)

For the UK, the 5 of 6 haplotypes (including H1-1-1, H1-2-1, H1-4-1, H1-4-2, and H1-4-3), which derived from H1 with previous 4 specific mutation sites (C241T, C3037T, C14408T, and A23403G) [6], accounted for 91.95% of the population (Table 4). H1-1-1 with only previous 4 specific mutation sites had almost disappeared in the UK by early 2021 (Fig 3). H1-2-1 with previous 4 specific mutation sites and the other 5 specific mutation sites (T445C, C6286T, C22227T, C26801G, and G29645T) appeared around July 21, 2020, became one of the major haplotypes circulating in the UK in early December 2020 [12], and gradually decreased, and there was only a very small population still circulating by late Feb, 2021 (Fig 3). While H1-4-1 with previous 4 specific mutation sites and another 17 specific mutation sites (C913T, C3267T, C5388A, C5986T, T6954C, C14676T, C15279T, T16176C, A23063T, C23604A, C23709T, T24506G, G24914C, C27972T, G28048T, A28111G, and C28977T) with mutation frequencies around 0.78, and H1-4-2 with one more mutation site (A17615G) compared with H1-4-1 showed a trend of increasing gradually since early December, 2020. And H1-4-1 and H1-4-2 had become the dominant epidemic haplotypes in the UK by early February, 2021 (Fig 3). Notably, the H1-4-1 and H1-4-2 haplotypes both had A23063T mutation causing the N501Y mutation on the S protein, and the N501Y mutation was almost completely linked with the other 16 mutation sites (C913T, C3267T, C5388A, C5986T, T6954C, C14676T, C15279T, T16176C, C23604A, C23709T, T24506G, G24914C, C27972T, G28048T, A28111G, and C28977T). Among the 17 sites, 11 caused amino acid changes, of which 5 mutation sites were located on the S protein (including N501Y, P681H, T716I, S982A, and D1118H) (Table 3). This may influence the epidemic traits of SARS-CoV-2 and the effectiveness of vaccines, especially mRNA vaccines.

**Figure 3:**
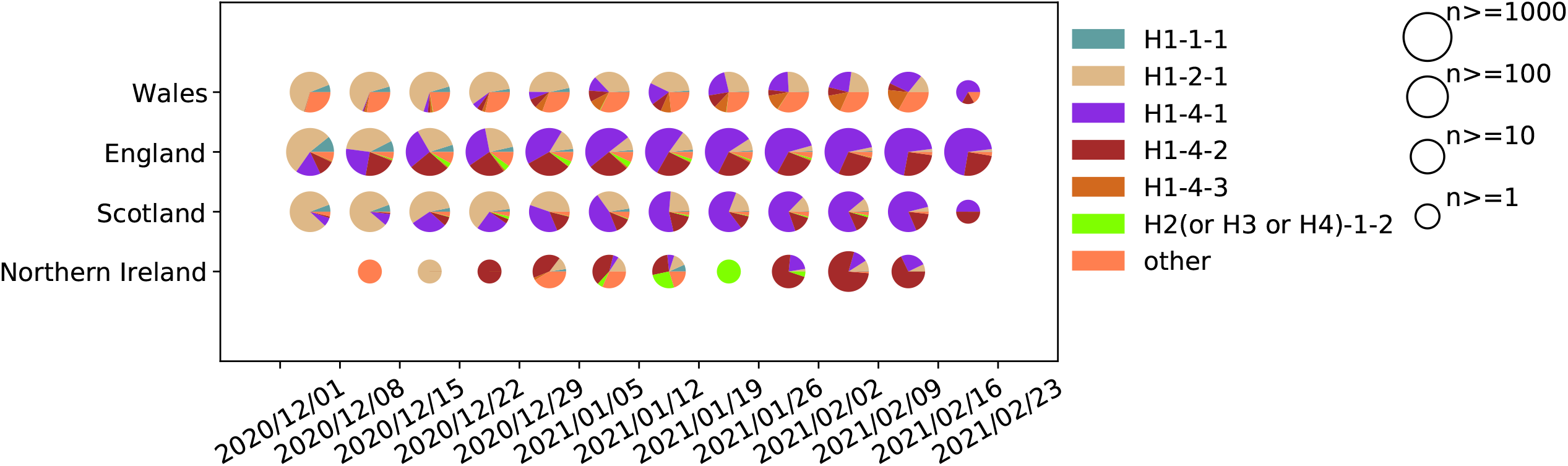
Epidemic trends of 6 haplotypes of 93,262 SARS-CoV-2 genomes from the UK

For Europe, the 5 haplotypes were the same as the 5 of 6 haplotypes of the UK (Table 4). Among the 5 haplotypes, 4 haplotypes (including H1-1-1, H1-2-1, H1-4-1, and H1-4-2) derived from H1 with previous 4 specific sites accounted for 84.67% of the population. And the epidemic trends of H1-1-1, H1-2-1, H1-4-1, and H1-4-2 were similar to those in the UK (Fig 4). That is, the H1-1-1 and H1-2-1 were gradually decreased, while the H1-4-1 and H1-4-2 were gradually increased.

**Figure 4:**
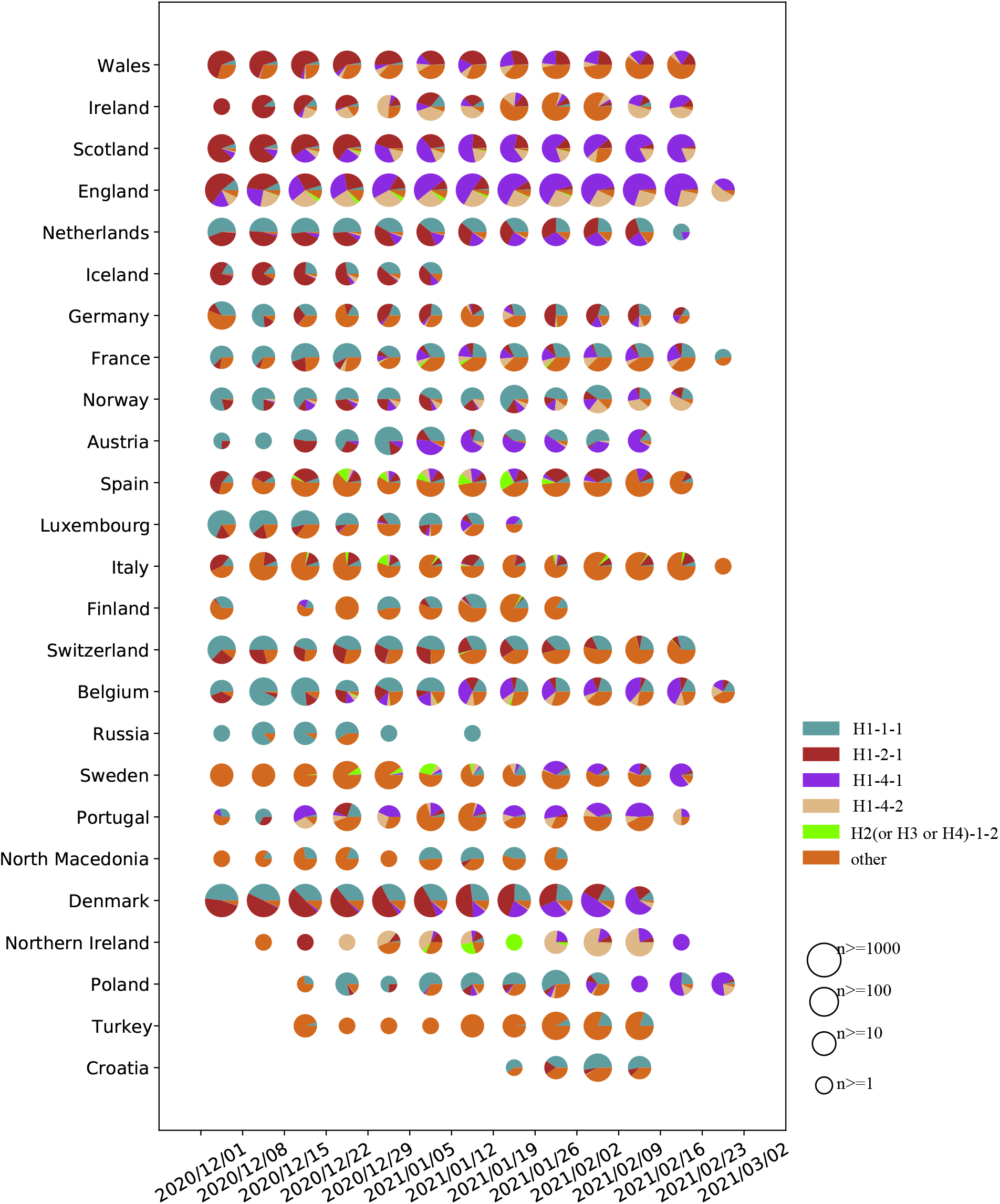
Epidemic trends of 5 haplotypes of 139,703 SARS-CoV-2 genomes from Europe. Countries or regions with a total number of genomes ≤ 100 were not shown in the figure.

### 3.3. Linkage and haplotype analysis of the 13 sites with a frequency ≥0.25 of SARS-CoV-2 in the USA

The detailed information for the 13 candidate key mutation sites screened from the USA was showed in Table 5. According to the linkage analysis, 21 haplotypes with a frequency ≥ 1% were found and accounted for 87.94% of SARS-CoV-2 population in the USA (Table 6), which showed some degree linked among the 13 candidate key mutation sites (Fig 2C). Among the 21 haplotypes, H1-1-1, H1-3-2, and H1-3-3, with a frequency >5%, all derived from H1 with previous 4 specific sites [6] (Table 6). H1-1-1 with previous 4 specific sites had a stable proportion (about 18%) between December 1, 2020 and February 28, 2021 in the USA (Fig 5). H1-3-2 and H1-3-3 were derived from H1-3 directly, and H1-3 derived from H1 directly with one more mutation site (G25563T) compared with H1 [6, 12]. H1-3-2 had previous 5 specific sites (C241T, C3037T, C14408T, A23403G, and G25563T) [12] and C1059T (Table 5, Table 6), which had a stable prevalent trend between December 01, 2020 and February 02, 2021 in the USA (Fig 5). H1-3-3 had previous 5 specific sites and 8 new missense mutation sites (C1059T, C10319T, A18424G, C21304T, G25907T, C27964T, C28472T, and C28869T) (Table 5, Table 6), which increased gradually between December 01, 2020 and February 02, 2021 in the USA (Fig 5). In general, the haplotype subgroup diversity in the USA is much more complicated than those of in the UK and Europe.

**Table 5.** The annotation of the 13 sites of SARS-CoV-2(USA) with a mutation frequency ≥25%

| Position | Ref | Alt | Frequency | Gene<br>Region | Mutation<br>Type | Protein<br>Changed | Codon<br>Changed | Impact |
| --- | --- | --- | --- | --- | --- | --- | --- | --- |
| 241 | C | T | 0.7720 | 5'UTR | upstream | NA | NA | MODIFIER |
| 1,059 | C | T | 0.6667 | orf1ab | missense | 265T>I | 794aCc>aTc | MODERATE |
| 3,037 | C | T | 0.9117 | orf1ab | synonymous | 924F | 2772ttC>ttT | LOW |
| 10,319 | C | T | 0.4435 | orf1ab | missense | 3352L>F | 10054Ctt>Ttt | MODERATE |
| 14,408 | C | T | 0.8505 | orf1ab | missense | 4715P>L | 14144cCt>cTt | MODERATE |
| 18,424 | A | G | 0.4625 | orf1ab | missense | 6054N>D | 18160Aat>Gat | MODERATE |
| 21,304 | C | T | 0.4593 | orf1ab | missense | 7014R>C | 21040Cgc>Tgc | MODERATE |
| 23,403 | A | G | 0.9454 | S | missense | 614D>G | 1841gAt>gGt | MODERATE |
| 25,563 | G | T | 0.6850 | ORF3a | missense | 57Q>H | 171caG>caT | MODERATE |
| 25,907 | G | T | 0.4846 | ORF3a | missense | 172G>V | 515gGt>gTt | MODERATE |
| 27,964 | C | T | 0.5072 | ORF8 | missense | 24S>L | 71tCa>tTa | MODERATE |
| 28,472 | C | T | 0.4827 | N | missense | 67P>S | 199Cct>Tct | MODERATE |
| 28,869 | C | T | 0.5029 | N | missense | 199P>L | 596cCa>cTa | MODERATE |

**Table 6.**
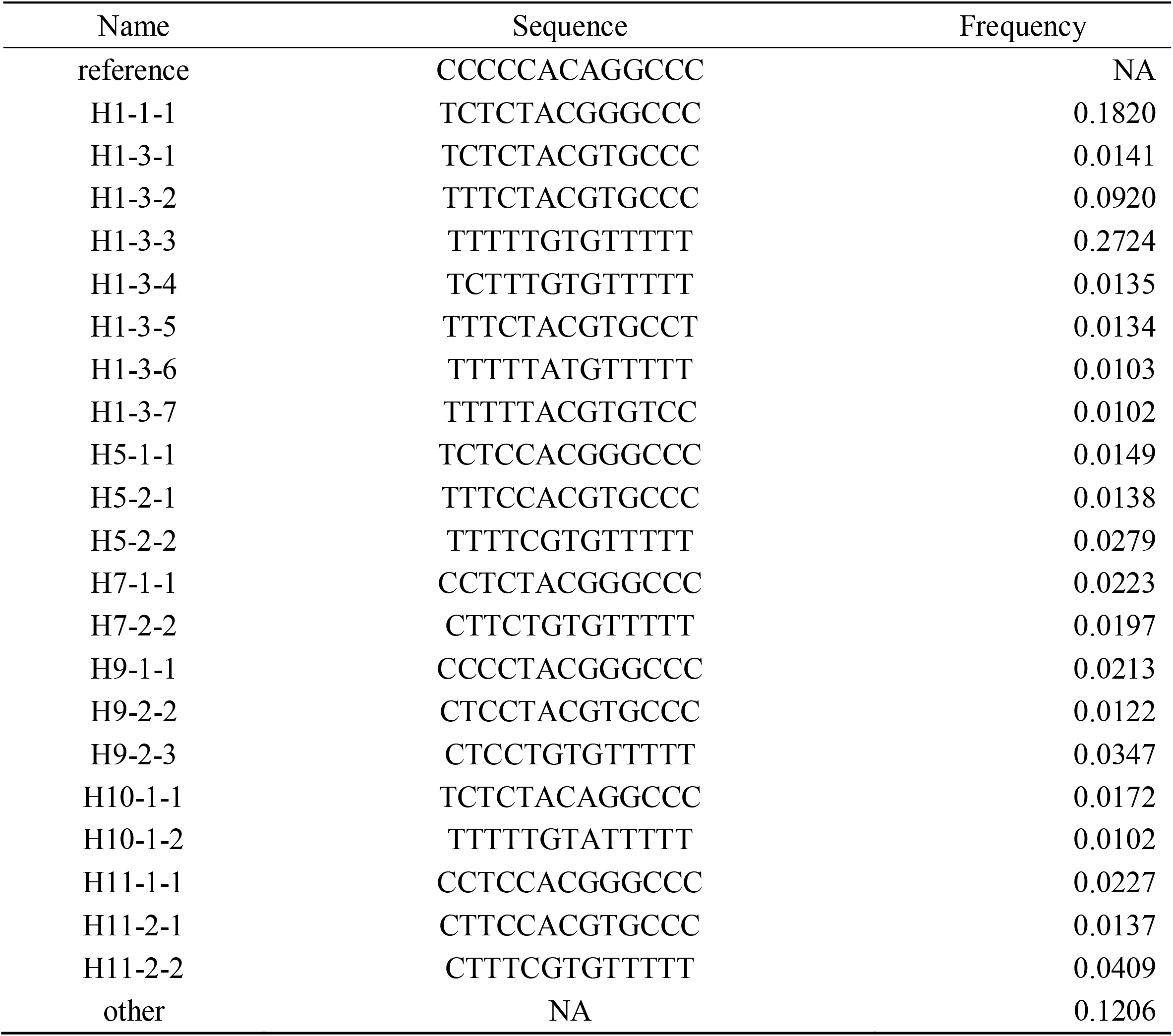
Haplotypes and their frequencies of the 13 sites of SARS-CoV-2(USA)

**Figure 5:**
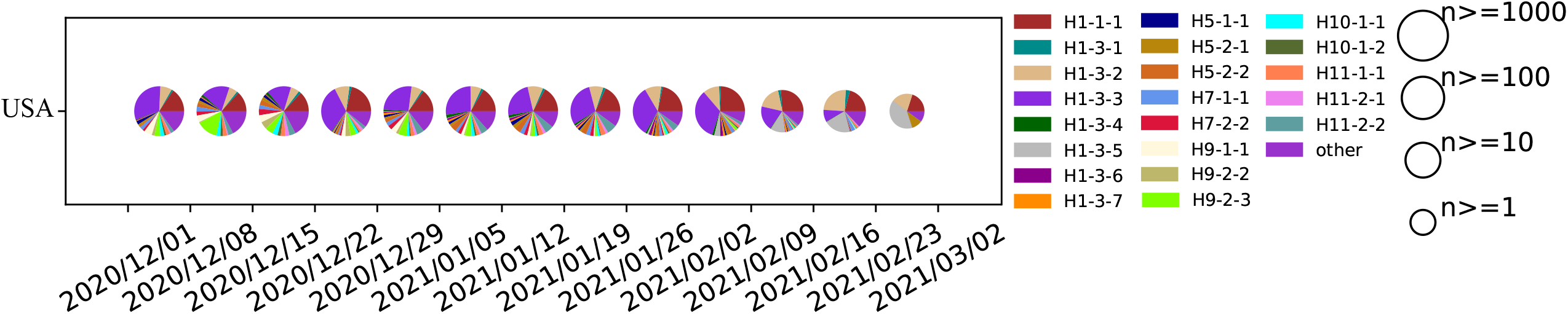
Epidemic trends of 21 haplotypes of 30,142 SARS-CoV-2 genomes from the USA

### 3.4. Linkage and haplotype analysis of 7 sites with a frequency ≥0.25 of HBV

The detailed information for the 7 candidate key mutation sites screened from HBV genomes was showed in Table 7. 5 of the 7 sites were missense mutations, including 356S>A (T192G), 444S>P (T456C), 807D>V (A1546T), 10R>K (G2337A) on P gene, and 331A>V (C659T) on the S gene (Table 7). These 5 mutations were all on the P gene or the overlapping part of the P gene and other genes. Linkage analysis and haplotype analysis were performed and found 24 haplotypes with a proportion ≥ 1%, of which there was not a major haplotype, indicating that the 7 sites of HBV had a low degree of linkage(Fig S1A, Table S1).

**Table 7.** The annotation of the 7 sites of HBV and 12 sites of HPV-16 with a mutation frequency ≥ 25%

| Virus | Position | Ref | Alt | Frequency | Gene Region<sup>1</sup> | Mutation Type | Protein Changed | Codon Changed |
| --- | --- | --- | --- | --- | --- | --- | --- | --- |
| HBV | 192 | T | G | 0.2804 | P, S | P: missense<br>S: synonymous | P: 356S>A<br>S: 175L | P: 356Tct>Gct<br>S: 175ctT>ctG |
|  | 456 | T | C | 0.2750 | P, S | P: missense<br>S: synonymous | P: 444S>P<br>S: 263Y | P: 444Tca>Cca<br>S: 263taT>taC |
|  | 659 | C | T | 0.5515 | P, S | P: synonymous<br>S: missense | P: 511S<br>S: 331A>V | P: 511agC>agT<br>S: 331gCc>gTc |
|  | 669 | C | T | 0.3205 | P, S | P: synonymous<br>S: synonymous | P: 515L<br>S: 334S | P: 515Ctg>Ttg<br>S: 334tcC>tcT |
|  | 1,546 | A | T | 0.4638 | P, X | P: missense<br>X: synonymous | P: 807D>V<br>X: 57G | P: 807gAc>gTc<br>X: 57ggA>ggT |
|  | 2,337 | G | A | 0.4863 | P, C | P: missense<br>C: synonymous | P: 10R>K<br>C: 174E | P: 10aGa>aAa<br>C: 174gaG>gaA |
|  | 2,479 | G | A | 0.4016 | P | P: synonymous | P: 57G | P: 57ggG>ggA |
| HPV-16 | 350 | T | G | 0.4508 | E6 | missense | 83L>V | 83Ttg>Gtg |
|  | 2,925 | A | G | 0.9157 | E2 | synonymous | 57Q | 57caA>caG |
|  | 3,409 | C | T | 0.5125 | E2, E4 | E2: missense<br>E4: synonymous | E2: 219P>S<br>E4: 26T | E2: 219Ccc>Tcc<br>E4: 26acC>acT |
|  | 3,977 | A | C | 0.5596 | E5 | missense | 39I>L | 39Ata>Cta |
|  | 4,040 | A | G | 0.6121 | E5 | missense | 60I>V | 60Ata>Gta |
|  | 4,226 | T | C | 0.4844 | Non-coding Region | NA | NA | NA |
|  | 4,363 | A | T | 0.9157 | L2 | missense | 43E>D | 43gaA>gaT |
|  | 4,936 | G | A | 0.8607 | L2 | synonymous | 234Q | 234caG>caA |
|  | 5,224 | A | C | 0.8192 | L2 | missense | 330L>F | 330ttA>ttC |
|  | 6,240 | C | G | 0.3415 | L1 | missense | 228H>D | 228Cat>Gat |
|  | 6,432 | A | G | 0.7019 | L1 | missense | 292T>A | 292Act>Gct |
|  | 7,191 | G | T | 0.6115 | Non-coding Region | NA | NA | NA |

<sup>1</sup>The HBV genome contains four genes: P gene, S gene, X gene, and C gene, some of which overlap partially

### 3.5. Linkage and haplotype analysis of 12 sites with a frequency ≥ 0.25 of HPV-16

The detailed information for the 12 candidate key mutation sites screened from HPV-16 genomes was showed in Table 7. Among them, 8 specific mutations were missense mutation, including 83L>V (T350G) on the E6 gene, 219P>S (C3409T) on the E2 gene, 39I>L (A3977C) and 60I>V (A4040G) on the E5 gene, 43E>D (A4363T) and 330L>F (A5224C) on the L2 gene, 228H>D (C6240G) and 292T>A (A6432G) on the L1 gene. Linkage analysis and haplotype analysis were performed on the 12 specific mutation sites and screened out 18 haplotypes with a proportion ≥1% (Table S2), and the 12 specific sites showed a low degree of linkage (Fig S1B). Among the 18 haplotypes, there were 5 major haplotypes with a frequency ≥4%, including H1, H2, H3, H4, and H5. The haplotype H2 had 5 specific mutation sites (A2925G, T4226C, A4363T, G4936A, and A5224C). H4 has 9 specific mutation sites (A2925G, C3409T, A3977C, A4040G, A4363T, G4936A, A5224C, A6432G, and G7191T), and H3 had one more mutation site (T350G) compared with H4, while H1 had two more mutation sites (T350G and T4226C) compared with H4. (Table 7, Table S2)

## Discussion

In this study, we developed a flexible tool to quickly monitor the candidate key mutations, haplotype subgroups, and epidemic trends for different viruses by using virus whole genome sequences, and analyzed a large number of SARS-CoV-2, HBV and HPV-16 genomes to show its functions, effectiveness and flexibility.

For the UK and Europe, we obtained the same 27 candidate key mutation sites, which could divide the SARS-CoV-2 population into 6 and 5 haplotypes, respectively. From the epidemic trend analysis, it showed that H1-4-1 and H1-4-2 with N501Y mutation on the S protein, which almost completely linked with the other 16 loci, had continued increasing from early December 2020 and became the dominant epidemic haplotypes in the United Kingdom and Europe by late February 2021. The B.1.1.7 mutant [23], corresponding to H1-4-1 and H1-4-2, has been reported that it has a more substantial transmission advantage based on several epidemiology researches [24, 25] and is greater in infectivity and adaptability [26]. Several studies have reported that the N501Y mutant reduces the neutralizing effect of the convalescent serum [27, 28], indicating that the N501Y variants may change neutralization sensitivity to reduce the effectiveness of the vaccine. Besides, the N501Y variants may reduce the effectiveness of antibodies [29]. Therefore, we should pay continuous attention to the N501Y mutant, which is almost completely linked with the other 16 loci.

For HPV-16 and HBV, we also screened out multiple specific sites which may be related to infectivity. For HPV-16, the T350G (83L>V) mutation we detected is the most common mutation on the E6 gene of HPV-16 [30-32]. Several studies have shown that the T350G mutant may cause persistent virus infection and further increase cancer risk [31-34]. It is reported that T350G variants can down-regulate the expression of E-cadherin, which is an adhesion protein that acts cell-cell adhesion. E-cadherin down-regulation can reduce the adhesion between cells, allowing infected cells to escape the host’s immune surveillance, and increase the risk of continued virus infection and the risk of cancer [33]. The C3410T mutation we detected, on the E2 gene of HPV-16, is also one of the common mutations of HPV-16 [35, 36]. Furthermore, The A2926G mutation we detected has been reported due to a reference genome sequencing error [37, 38]. For HBV, the C659T mutation, which causes A331V mutation on S gene, is reported to be associated with increasing the efficiency of HBV replication [39].

Due to the continuous mutations and evolution of viruses, it should be carefully considered whether the new mutations have an influence on developing and updating vaccines. AutoVEM2 provides a fast and reliable process of continuously monitoring candidate key mutations and epidemic trends of these mutations. Through AutoVEM2, we have analyzed a large number of SARS-CoV-2, HBV, and HPV-16 genomes and obtained some candidate key mutation sites fast and effectively. Among them, some mutations, such as D614G and N501Y of SARS-CoV-2, T350G of HBV, and C659T of HVP-16, have been proved to play an important role in the viruses, indicating the reliability and effectiveness of AutoVEM2. In total, we developed a flexible automatic tool for monitoring candidate key mutations and epidemic trends for any virus. It can be used in the study of mutations and epidemic trends analysis of existing viruses, and can be also used in analyzing the virus that may appear in the future. Our integrated analysis method and tool could become a standard process for virus mutation and epidemic trend analysis based on genome sequences in the future.

## Authors’ contributions

BX developed the tool, carried out the data analysis, and wrote the manuscript. SL collected the data and wrote the manuscript. WL collected the data. DW, YB, YQ, RL, and LH revised the manuscript. HD conceived and supervised the study and revised the manuscript.

## Availability

The developed AutoVEM2 software has been shared on the website (https://github.com/Dulab2020/AutoVEM2) and can be freely available.

## Data availability

All data relevant to the study are included in the article or uploaded as supplementary information.

## Ethical Approval

Not required.

## Tables and Figures Legends

**Figure S1**: Linkage analysis of candidate key mutation sites of HBV and HPV-16.

A. Linkage analysis of 7 candidate key mutation sites of HBV;

B. Linkage analysis of 12 candidate key mutation sites of HPV-16

**Table S1**: Haplotypes and their frequencies of the 7 sites of HBV

**Table S2**: Haplotypes and their frequencies of the 12 sites of HPV-16

