## Supplementary figures and images for "AutoVEM2: a flexible automated tool to analyze candidate key mutations and epidemic trends for virus"

### Fig S1A, Fig S1B

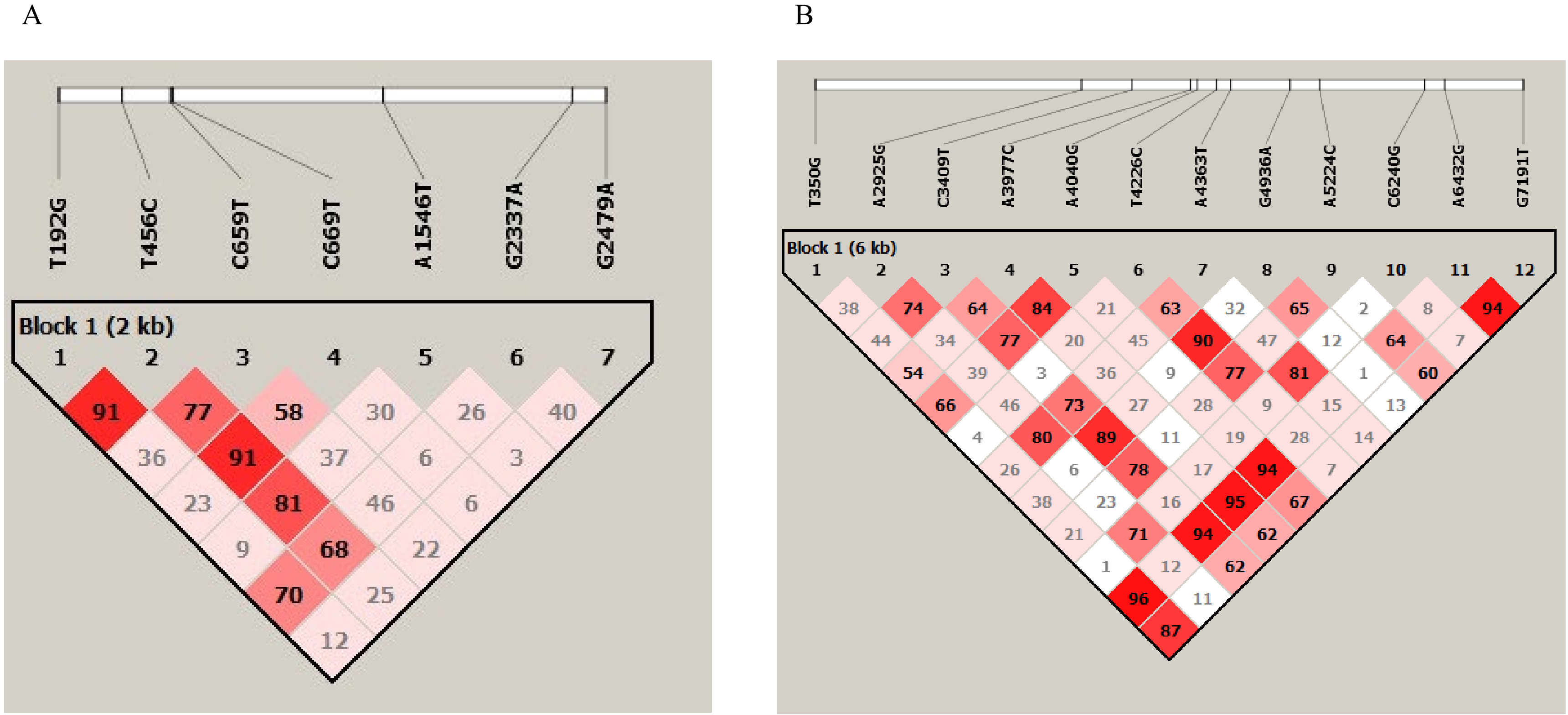
